# Fbxo45 binds SPRY motifs in the extracellular domain of N-cadherin and regulates neuron migration during brain development

**DOI:** 10.1101/828426

**Authors:** Youn Na, Elisa Calvo-Jiménez, Elif Kon, Hong Cao, Yves Jossin, Jonathan A. Cooper

**Author notes:** YN; Seattle Genetics, Seattle, WA, HC; Institutes of Brain Science, State Key Laboratory of Medical Neurobiology and Collaborative Innovation Center for Brain Science, Fudan University, 200032, China. Contributing authors (Yves Jossin, Jonathan A. Cooper).

## Abstract

Several events during normal development of the mammalian neocortex depend on N-cadherin, including the radial migration of immature projection neurons into the cortical plate. Remarkably, radial migration requires the N-cadherin extracellular domain but not N-cadherin-dependent homophilic cell-cell adhesion, suggesting that other N-cadherin-binding proteins may be involved. We used proximity ligation and affinity purification proteomics to identify N-cadherin-binding proteins. Both screens detected MycBP2 and SPRY-domain protein Fbxo45, two components of an intracellular E3 ubiquitin ligase. Fbxo45 appears to be secreted by a non-classical mechanism, not involving a signal peptide and not requiring endoplasmic reticulum to Golgi transport. Fbxo45 binding requires N-cadherin SPRY motifs that are not involved in cell-cell adhesion. SPRY-mutant N-cadherin does not support radial migration in vivo. Radial migration was similarly inhibited when Fbxo45 expression was suppressed. The results suggest that projection neuron migration requires both Fbxo45 and binding of Fbxo45 or another protein to SPRY motifs in the extracellular domain of N-cadherin.

## INTRODUCTION

The complex layered structure of the mammalian neocortex arises through the coordinated generation, specification, migration and connection of different types of neurons (1-3). Projection neurons are born and specified in the pallial ventricular zone (VZ) but journey long distances before undergoing terminal differentiation. The journey occurs in stages. First, newborn neurons enter the subventricular zone/intermediate zone, become multipolar, and migrate randomly (4). At the top of the intermediate zone, signals induce multipolar neurons to migrate radially outwards into the cortical plate (CP), becoming bipolar and attaching to radial glia as they move. They then pass beyond earlier-born neurons and reach the top of the CP, where they undergo terminal translocation, stop migrating, and differentiate. This pattern of migration gives rise to the classic “inside-out” lamination of the neocortex, with first-born neurons positioned inside later ones. Genetic disruption of neuron migration is associated with neurodevelopmental disorders including lissencephaly, epilepsy and schizophrenia.

Classical cadherins, including neuronal (N-)cadherin (NCad) and epithelial (E-)cadherin (ECad), are calcium-dependent cell-cell adhesion molecules (5). NCad is important for cell-cell adhesion in the neuroepithelium of the VZ (6, 7). NCad also regulates projection neuron migration at two stages: first when neurons enter the CP (8-10), and second, during terminal translocation (11-13). NCad is upregulated on the cell surface at both these stages in response to an extracellular signal, Reelin. Reelin stimulates signaling pathways involving Src kinases, Dab1, PI3’ kinase, Crk/CrkL, C3G, and the small GTPases, Rap1 and RalA (14-21). These pathways increase NCad surface expression, in collaboration with Rab GTPase and Drebin-like vesicular trafficking (9, 11, 22, 23). How NCad regulates neuron migration is unclear, however.

Because cadherins are best known for cell-cell adhesion (5), neuron migration may require homophilic binding of NCad on migrating neurons to NCad on other neurons, axons, or radial glia. Indeed, cultured neurons will polarize towards an external source of NCad (24). NCad may also be involved in attaching bipolar neurons in the CP to radial glia (25). However, our recent work revealed that a NCad^W161A^ mutant, which cannot form “strand-swap” trans homodimers or support cell-cell adhesion (26-29), can support neuron migration into the CP (30). We further found that CP entry requires NCad binding to, and activating, fibroblast growth factor receptors (FGFRs) in cis (on the same cell) (30). ECad does not support CP entry even though it binds FGFR. Mechanistically, NCad but not ECad protected FGFR from degradation, and the first two of five extracellular calcium-binding domains on NCad (EC1 and EC2) were critical. These results leave open the possibility that additional proteins binding NCad but not ECad may regulate neuron migration.

Fbxo45 (F box/SPRY domain-containing protein 1) is a little-studied protein that is highly expressed in the nervous system and required for cortical lamination, axonal outgrowth and synaptic connectivity (31-33). Most F-box proteins bind Skp1, Cul1 and Rbx1 to form a SCF (Skp1-Cul1-F-box) E3 ubiquitin ligase complex. Fbxo45 is atypical in that it does not bind Cul1 or Rbx1 and instead associates with MycBP2/PAM (Myc binding protein 2/protein associated with Myc), forming a Fbxo45-Skp1-MycBP2 complex that has E3 ligase activity in vitro (32). The SPRY domain of Fbxo45 potentially interacts with substrates. Curiously, NCad was detected in a Fbxo45 interaction screen (34). Furthermore, knockdown of Fbxo45 decreased NCad expression and impaired differentiation of neuronal stem cells (34), suggesting that Fbxo45 interaction with NCad is involved in brain development.

Here we set out to identify secreted proteins that interact with the ectodomain of NCad and may regulate radial polarization of multipolar neurons. Two different unbiased proteomics approaches detected Fbxo45 and MycBP2 as major binding partners for the extracellular domain of NCad. We found that the Fbxo45 SPRY domain binds to SPRY motifs in the EC1 region of NCad but not ECad. Mutation of these motifs does not inhibit cell-cell adhesion but does inhibit neuron migration into the cortical plate in vivo. Fbxo45 appears to be secreted by an unconventional mechanism independent of a signal peptide. Cell autonomous knockdown of Fbxo45 inhibited entry into the cortical plate. These results suggest that Fbxo45 is secreted and required for normal neuron migration during brain development, potentially by binding the extracellular domain of NCad.

## RESULTS

### BioID identifies Fbxo45 and MycBP2 as extracellular NCad-interacting neuronal proteins

We used two proteomics approaches to identify NCad binding proteins: proximity-dependent biotin identification (BioID) (35), in which mutant BirA^R118G^ (BirA*) transfers biotin onto nearby amino group, and affinity purification followed by mass spectrometry. For BioID, we fused a Myc-tagged BirA^R118G^ (BirA*) in the extracellular domain of HA-tagged NCad (NCad-HA), expressed the fusion protein on HEK293T cells, and cultured the cells with primary rat embryonic cortical neurons. The mixed cell population was incubated with biotin and ATP and biotinylated proteins were isolated and identified. The biotinyl-AMP generated by BirA* has a working distance ∼10 nm (36), which is about half the length spanned by the five extracellular cadherin (EC) repeats (37) (diagrammed in Figure 1A). To detect proteins that might interact with either end of the NCad ectodomain, we inserted Myc-BirA* either in the middle of EC2 (N2-BirA*) or between EC5 and the transmembrane domain (TM) (N5-BirA*) (Figure 1A). Insertion of small protein domains at these sites does not to interfere with adhesion (37). As a control, the EC1-5 region of N5-BirA* was replaced with EC1-5 of ECad, to create E5-BirA*. All fusion proteins appeared to traffic normally to the cell surface (Figure 1A). In addition, NCad was biotinylated when cells co-transfected with NCad and N5-BirA* were labeled with ATP and biotin (Figure 1B). These results suggest that N5-BirA* co-clusters with NCad at the surface.

**Figure 1.**
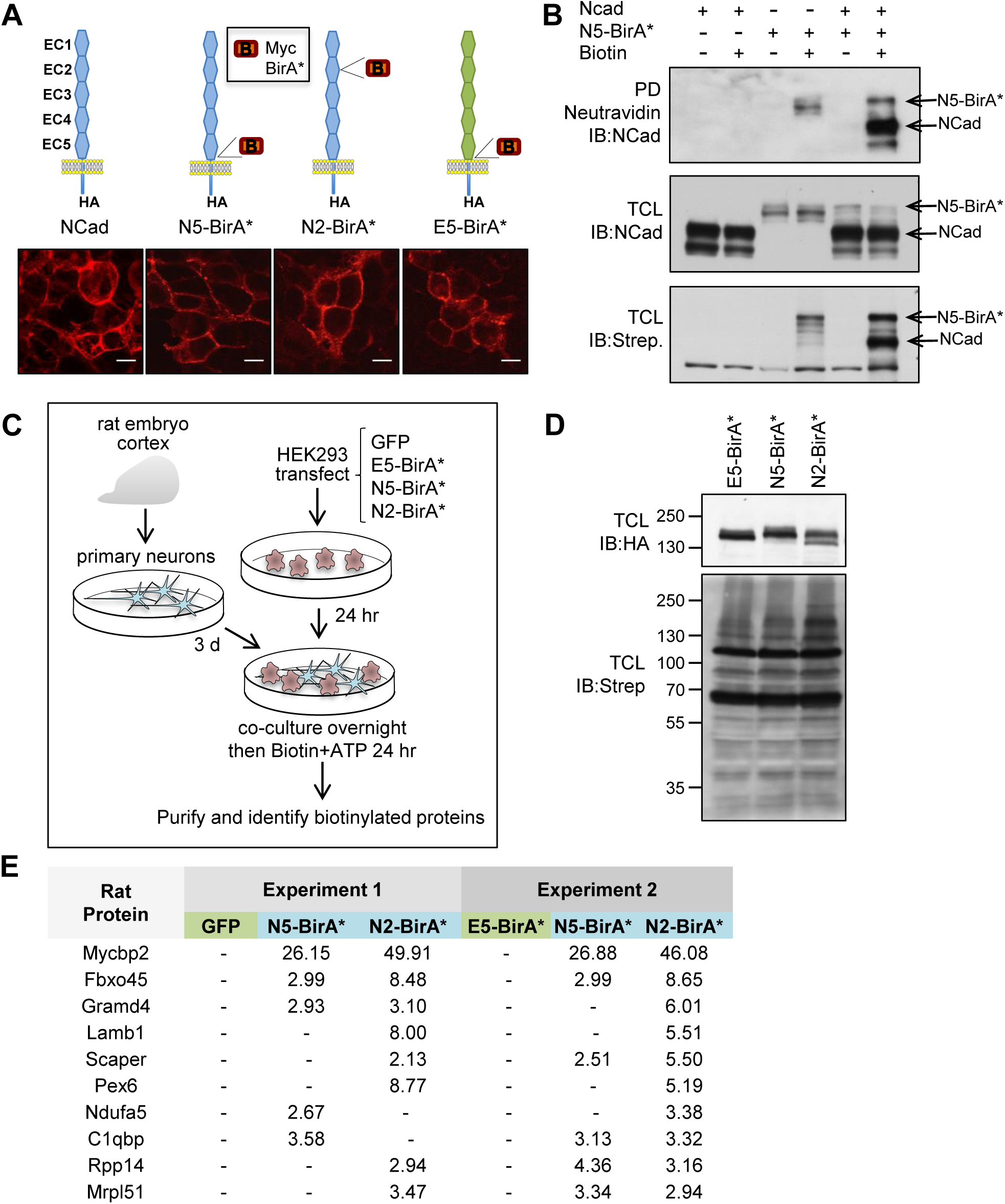
Characterization of BirA*fusion proteins and BioID screen for NCad *trans*-interacting proteins. (A) Structure and localization of cadherin fusion proteins. Upper: cadherin expression constructs. NCad, ECad and myc-BirA* are marked blue, green and red. All constructs have a C-terminal HA tag. Bottom: Representative immunofluorescence images of the respective constructs expressed in HEK293T cells. Scale bar: 10 μm. (B) Biotinylation of NCad by N5-BirA*. HEK293T cells were transfected with NCad or N5-BirA* separately or together. PD: Pull-down with Neutravidin beads followed by blot with NCad antibody. TCL: total cell lysates probed with NCad antibody or streptavidin-HRP. (C) Strategy for BioID screen. HEK293T cells were transfected with the indicated constructs then co-cultured with DIV3 rat cortical neurons in the presence of biotin and ATP. Cell lysates were purified using Streptavidin beads. (D) Total cell lysates were analyzed by Western blotting with anti-HA to detect the fusion proteins and Streptavidin-HRP to detect biotinylated proteins. (E) Samples were analyzed by on-bead trypsin digestion and LC-MS/MS. Highest abundance rat neuron proteins (peptide spectrum matches) biotinylated in the presence of N5-BirA*, N2-BirA* and negative controls GFP (experiment 1) or E5-BirA* (experiment 2). -, undetected.

To identify neuronal proteins that interact with the NCad extracellular domain, HEK293T cells were transiently transfected with N5-BirA*, N2-BirA*, or negative control plasmids GFP (experiment 1) or E5-BirA* (experiment 2). The transfected cells were cultured with rat embryonic cortical neurons and labeled with biotin and ATP (Figure 1C). Cell lysates were collected and samples were analyzed for protein expression (top, Figure 1D) and biotinylation (bottom, Figure 1D). In all cases there was extensive biotinylation of many proteins. The remaining sample was purified using Streptavidin beads and digested with trypsin before LC-MS/MS analysis. The identified peptides were searched against the rat protein database to distinguish rat neuron proteins from human HEK293T proteins (Figure 1E and supplementary tables 1 and 2). MycBP2 and Fbxo45 were the highest-ranked rat neuron proteins in both experiments (Figure 1E and Supplementary Table 1). MycBP2 also was the highest-ranked human protein in both experiments (Supplementary Table 2). The results suggest that Fbxo45 and MycBP2 are major proteins expressed by rat neurons that interact with the ectodomain of NCad. However, eight out of the top 10 proteins, including Fbxo45 and MycBP2, are predicted intracellular proteins, so it is not clear whether they were released by cell death or by an active process.

### Affinity purification of neuronal proteins that bind to NCad

As an independent approach to detect proteins that might regulate neuron migration, we affinity purified proteins from embryonic mouse brain homogenate using recombinant NCad^W161A^ ectodomain, which does not form strand-swap dimers, as bait. The ectodomain was fused to the human immunoglobulin constant region (Fc), expressed in HEK293T cells, and purified from the culture supernatant using a Protein A/G (PAG) column. An ECad-Fc fusion protein was prepared similarly as a negative control. Neuronal proteins were then purified as illustrated in Figure 2A. Mouse embryonic forebrains were homogenized, insoluble material was removed by centrifugation, and the supernatant passed over PAG. Flow through from the PAG column was then passed over ECad-Fc, to remove proteins that bind to ECad. The unbound material was then passed over NCad^W161A^-Fc, to select proteins that bind to NCad^W161A^. All three columns were washed and eluted at low pH. As expected, PAG retained large amounts of immunoglobulin heavy and light chains (IgH and IgL, lane 7, Figure 2B) while ECad-Fc and NCad^W161A^-Fc eluates contained major amounts of ∼110 kDa proteins corresponding to the cadherin-Fc fusions (lanes 9 and 11, Figure 2B). Following preparative SDS polyacrylamide electrophoresis, ECad-Fc and NCad^W161A^-Fc proteins running above and below the ∼110 KDa cadherin-Fc bands were trypsinized and identified by LC-MS/MS. The 10 most abundant proteins that bound to NCad^W161A^ but not to ECad were predicted intracellular, and presumably were released from dead or broken cells (Figure 2C, Supplementary Table 3). MycBP2 and Fbxo45 were the major proteins. Skp1 was also detected. Our results are consistent with NCad binding to an Fbxo45-Skp1-MycBP2 complex that is secreted or released from cells.

**Figure 2.**
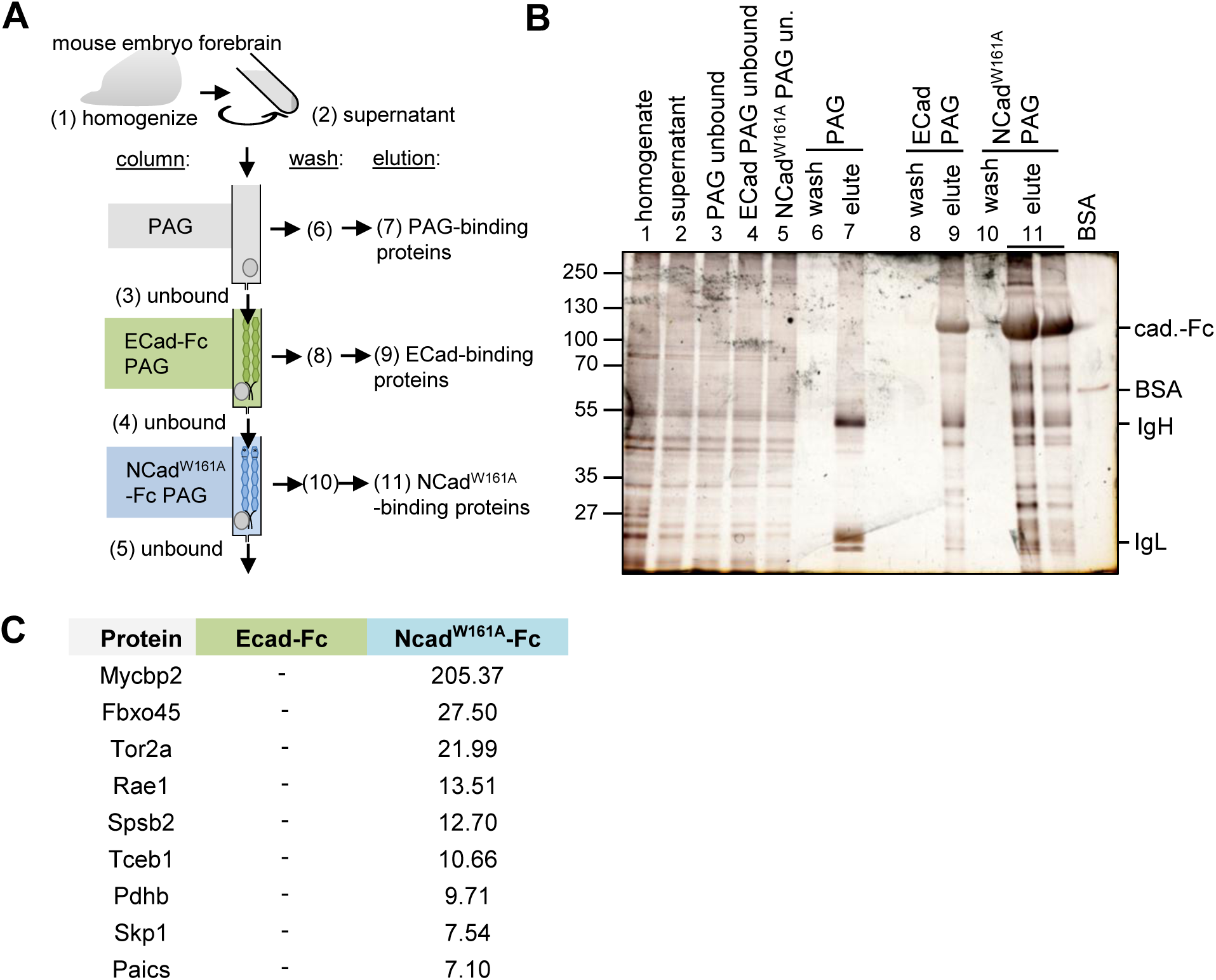
Affinity purification screen for brain proteins that bind the NCad but not ECad extracellular domain. (A) Strategy for affinity purification of NCad^W161A^ -specific embryonic brain proteins. Numbers in parentheses correspond to samples analyzed in (B). PAG: Protein A/G Sepharose. (B) Silver stained SDS polyacrylamide gel of samples from the purification. (C) Highest abundance proteins detected binding to NCad^W161A^ and ECad (peptide-spectrum matches). Fractions 9 and 11 were separated by preparative SDS polyacrylamide gel electrophoresis and regions above and below the cadherin-Fc band excised. Proteins in these regions were analyzed by in-gel trypsin digestion and identified by LC-MS/MS.

### The EC1-2 region of NCad interacts with the SPRY domain of Fbxo45

To confirm that Fbxo45 binds to NCad but not ECad, Fbxo45 was tagged with T7 and co-transfected with HA-tagged cadherins. As expected, Fbxo45 co-immunoprecipitated with NCad but not ECad (Figure 3A). When the ectodomain of NCad was deleted (ΔEC) or the EC1-2 domains of NCad were switched to ECad (EN), Fbxo45 binding was inhibited (Figure 3B). Moreover, NCad EC1-2 was sufficient to bind Fbxo45 (Figure 3C). Taken together, our data suggest that the EC1-2 domains of NCad but not ECad are necessary and sufficient to bind Fbxo45. This finding is consistent with our observation that Fbxo45 was more efficiently biotinylated if BirA* was fused into EC2 than into the juxtamembrane region (compare N2-BirA* and N5-BirA*, Table 1), and with a previous report (34).

**Figure 3.**
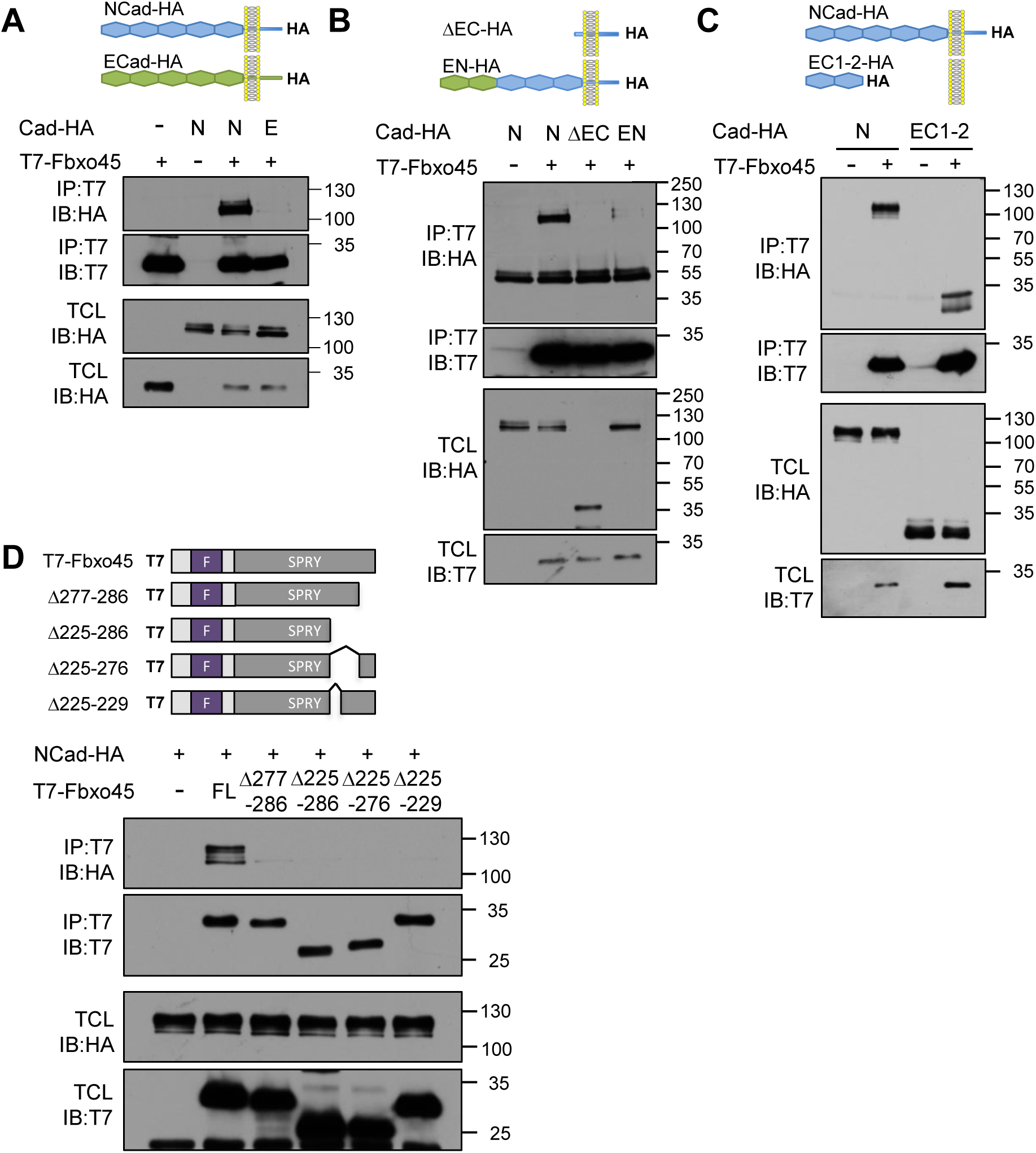
NCad EC1-2 interacts with SPRY domain of Fbxo45. (A-C) Various forms of NCad-HA were co-transfected with T7-Fbxo45 into HeLa cells. Cell lysates were immunoprecipitated with T7 antibody and samples of cell lysates (TCL) and immunoprecipitates (IP) analyzed by Western blotting. (A) Fbxo45 interacts with NCad, but not ECad. (B) NCad EC1-2 is necessary to interact with Fbxo45. (C) NCad EC1-2 is sufficient to interact with Fbxo45. (D) SPRY domain of Fbxo45 is necessary to interact with NCad. All deletion mutations in the SPRY domain disrupt Fbxo45 binding to NCad.

We next investigated the NCad-binding region of Fbxo45. Fbxo45 contains an F-box domain (residues 33-83) that binds Skp1 and a SPRY domain (92-283) that binds ubiquitin ligase substrates (32). We deleted different regions of the SPRY domain and assayed binding to NCad (Figure 3D). Remarkably, none of the mutants tested was able to bind NCad, even when we deleted as little as 10 residues from the C terminus (mutant 276) or five residues between 225 and 229 (Δ225-229). These regions correspond to β strands 11 and 15 in the predicted structure of the Fbxo45 SPRY domain (38). Our results implicate the folded structure of the Fbxo45 SPRY domain in binding to NCad.

### SPRY-binding motifs in NCad

We reasoned that the Fbxo45 SPRY domain may bind to one or more of three canonical SPRY motifs ([DE][IL]NXN (39)) in NCad EC1-2 that are not conserved in ECad (Figure 4A). Chung et al (34) previously reported that deletion of motifs 2 and 3 abolished Fbxo45 binding. However, motifs 2 and 3 chelate calcium ions in the folded structure (27, 40, 41). Deleting these motifs might cause mis-folding and indirectly disrupt Fbxo45 binding. We therefore made conservative mutations, substituting SPRY motifs 1, 2 or 3 with the corresponding sequences from ECad, generating chimeras NC1, NC2 and NC3. These chimeras were then tested for co-immunoprecipitation with Fbxo45 (Figure 4B). Substitution of motif 1 (D242S, I243S, N246E and Q247A) almost completely inhibited interaction, while substitution of motif 2 (M261Q) or motif 3 (V376I and P380A) was less inhibitory. Substituting motifs 1 and 2 (NC12), or 1, 2 and 3 (NC123), abolished binding (Figure 4B). However, NC123 still bound NCad (Figure 4C) and was expressed on the surface of transfected cells, co-localizing with endogenous epidermal growth factor receptor (EGFR) (Figure 4D).

**Figure 4.**
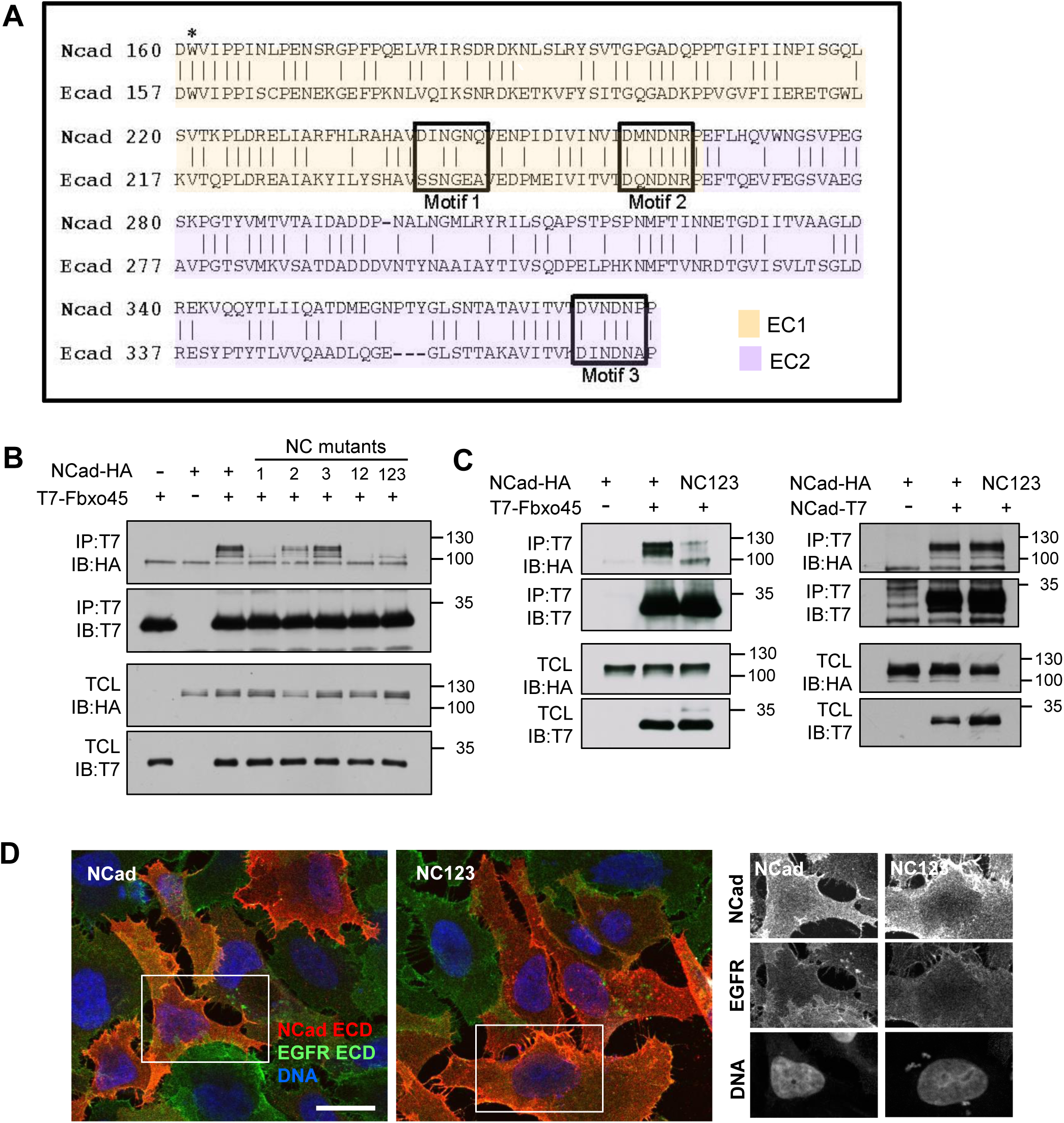
SPRY motifs in NCad bind Fbxo45. (A) Alignment of mouse NCad and ECad EC1-2 sequences. The alignments start with the first residues of the cleaved cadherins, while the numbering corresponds to the primary translation products. Motifs 1-3: three matches to the SPRY recognition motif ([DE][IL]NXN) in NCad. Note that motifs 2 and 3 also bind calcium. * indicates W161, which is critical for the strand-swapped trans dimer. (B) Identification of the Fbxo45 binding site. NCad-HA chimeras containing ECad residues from motifs 1, 2 or 3, were co-transfected into HeLa cells with T7-Fbxo45. Cell lysates were immunoprecipitated using T7-antibody. (C) NC123 fails to bind Fbxo45 but still binds to NCad. (D) NC123 traffics to the cell surface. HeLa cells were transfected to express NCad-HA or NC123-HA, fixed, and immunostained with antibodies to the extracellular domain of NCad and EGFR. White boxes correspond to individual channels shown at right. Scale bar: 20 μm.

To test whether the Fbxo45-binding sites were required for homophilic adhesion, CHO-K1 cells, which lack cadherins (42), were transfected with GFP and wildtype or mutant NCad. Transfected cells were allowed to aggregate in the presence or absence of calcium. Calcium-dependent aggregation was stimulated by wildtype NCad or NC123 but not by NCad^W161A^ (Figure 5A and B). Moreover, mCherry-labeled cells expressing NC123 but not NCad^W161A^ aggregated with GFP-labeled wildtype NCad cells in the presence of calcium (Figure 5C). Thus NC123 binds to NCad but not Fbxo45 while NCad^W161A^ binds to Fbxo45 but not NCad.

**Figure 5.**
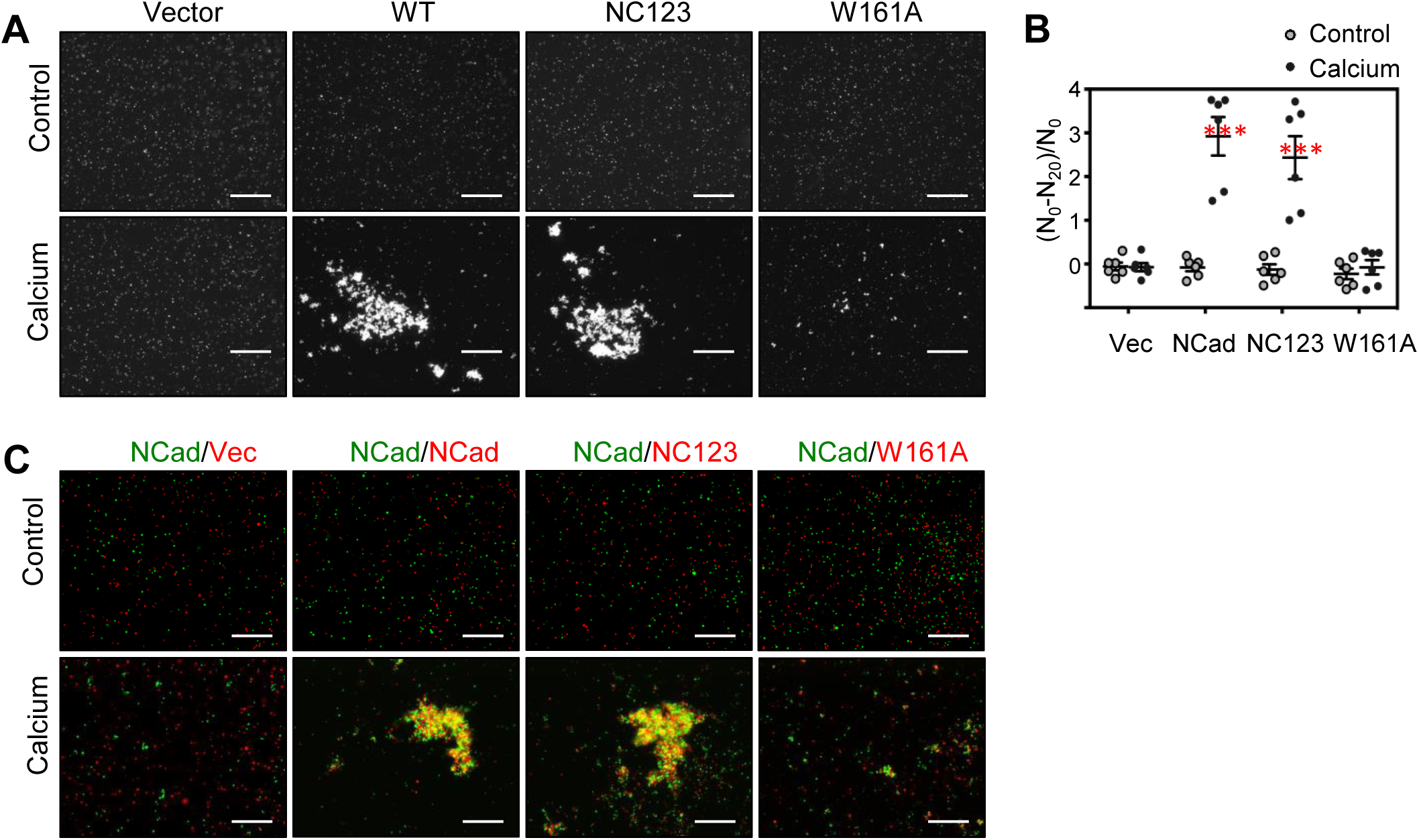
SPRY motif mutant NC123 supports calcium-dependent cell-cell interaction. (A) Aggregation assay. The indicated constructs were transfected into CHO-K1 cells together with GFP as a marker for transfected cells. Cell suspensions were allowed to aggregate in the absence or presence of calcium. Scale bar: 800 μm. (B) Quantification of replicate experiments. N_0_: particle number at time 0 min, N_20_: particle number at time 20 min. n = 6. ***p <0.001. (C) Aggregation assay with two different cell populations to investigate trans-interaction between NCad wildtype and mutants. Green cells, co-expressing NCad wildtype and GFP, were mixed with red cells, co-expressing NCad wildtype or mutants and mCherry. Scale bar: 400 μm.

### Fbxo45 reaches the cell surface through a non-classical secretion pathway

The binding of Fbxo45 to EC1-2 of NCad raises the question of how Fbxo45 reaches the cell exterior. Transfected cells released T7-Fbxo45 or Fbxo45-T7 but not tubulin, suggesting that Fbxo45 is actively secreted and not released from broken cells (Figure 6A and data not shown). While Fbxo45 lacks an N-terminal endoplasmic reticulum (ER) translocation signal for conventional secretion (43), some proteins that lack signal peptides are secreted by unconventional pathways (44-47). To test whether Fbxo45 is secreted by an unconventional pathway, we used Brefeldin A (BFA), which inhibits conventional but not unconventional secretion (48). BFA slightly increased secretion of T7-Fbxo45 but inhibited secretion of a similar-sized N-terminal signal sequence protein, EC1-T7 (Figure 6A). This suggests that Fbxo45 is secreted by a BFA-insensitive unconventional route.

**Figure 6.**
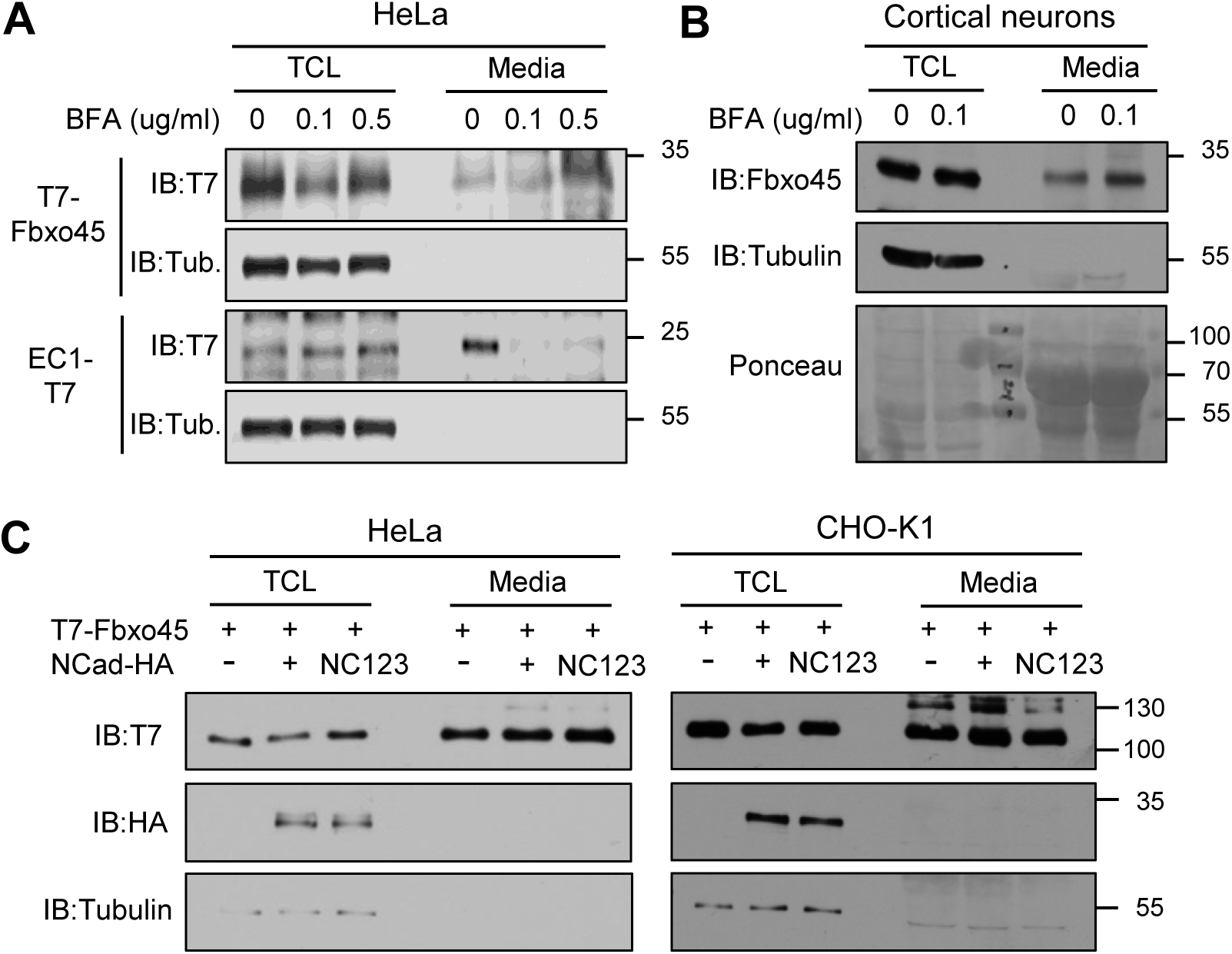
Fbxo45 is secreted by an unconventional pathway. (A) T7-Fbxo45 secretion is secreted by a Brefeldin A (BFA)-insensitive route. HeLa cells were transfected with NCad EC1-T7 or T7-Fbxo45 and incubated with various concentrations of BFA in serum-free media for 24 hours. Cell lysates (TCL) and concentrated media (Media) were harvested. 0.5 % of the TCL and 15 % of the Media were analyzed by Western blotting. Tub.: tubulin control for cell lysis. (B) Mouse primary cortical neurons secrete Fbxo45 by an unconventional pathway. DIV 2 cultures were incubated with serum-free Neurobasal media containing 0 or 0.1 μg/ml of BFA for 20-24 hours. Media and TCL were subject to Western blotting for Fbxo45 and tubulin. Ponceau staining shows equal loading. (C) Co-expression of NCad or NC123 with Fbxo45 does not affect Fbxo45 secretion by HeLa or CHO-K1 cells.

To test whether untagged, endogenous Fbxo45 is secreted, primary mouse cortical neurons were incubated with serum-free Neurobasal media with or without BFA for 20-24 hrs. Media were collected and Fbxo45 detected with a specific antibody (34). Fbxo45 but not tubulin was detected in the neuron culture media, and secretion was increased by BFA, suggesting unconventional secretion (Figure 6B).

NCad could potentially bind Fbxo45 in an intracellular vesicle and help ferry Fbxo45 to the surface. Therefore, we tested whether co-expression of NCad with Fbxo45 would stimulate Fbxo45 secretion. Neither NCad nor NC123, which does not bind Fbxo45, affected secretion of Fbxo45 from HeLa or cadherin-deficient CHO-K1 cells (Figure 6C). Taken together these results suggest that Fbxo45 is secreted by an unconventional, NCad-independent pathway.

### Neuron migration into the cortical plate requires Fbxo45 and NCad SPRY motifs

Since ECad EC1-2 cannot substitute for NCad EC1-2 for either Fbxo45 binding or neuron migration (30), Fbxo45 may regulate neuron migration. We tested whether Fbxo45 regulates neuron migration by in utero electroporation of GFP and Fbxo45 shRNA at embryonic day (E) 14.5, and measuring the positions of GFP-expressing neurons three days later. Under these conditions, Fbxo45 shRNA (shFbxo45) significantly inhibited movement of neurons from the multipolar migration zone (MMZ) to radial migration zone (RMZ) (Figure 7A and C, P=0.0003). A similar arrest was observed with another shRNA targeting a different sequence in Fbxo45 (not shown). Immunofluorescence of cortical sections two days after electroporation (E16.5), when neuron migration is not yet delayed, showed no effects on proliferation (Ki67), differentiation (Sox2, Tbr2 or Tuj1), apoptosis (cleaved Capase3) or the radial glia fibers (Nestin: Figure 7D). These results suggest that Fbxo45 is needed for neuron entry into the cortical plate, but do not distinguish whether Fbxo45 is required inside or outside migrating neurons.

**Figure 7.**
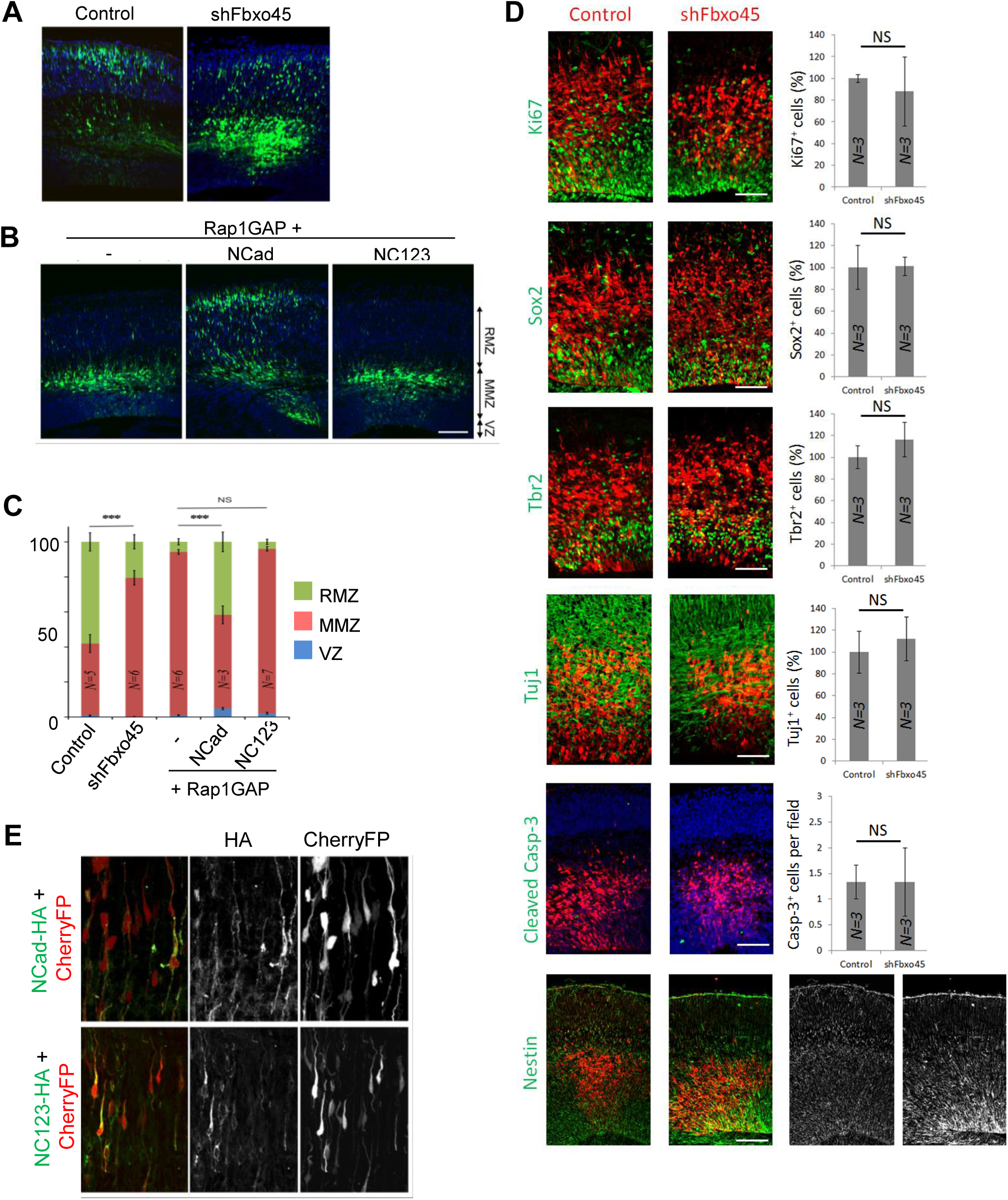
Fbxo45 and NCad SPRY-motifs are required for neuron entry into radial migration zone in vivo. (A) Fbxo45 shRNA inhibits neuron migration from the multipolar migration zone (MMZ) to the radial migration zone (RMZ) with little effect on the ventricular zone (VZ). E14.5 mouse embryos were microinjected and electroporated in utero with plasmids expressing GFP alone (control) or with Fbxo45 shRNA. Three days later, sections were prepared and the positions of GFP-expressing neurons were recorded. Representative sections are shown. (B) NC123 does not support migration of Rap1GAP-expressing neurons into the RMZ. E14.5 mouse embryos were microinjected and electroporated in utero with plasmids expressing GFP and Rap1GAP together with vector, NCadHA or NC123HA. Analysis as in (A). (C) Quantification of (A) and (B). Percentage of GFP-expressing neurons in the VZ, MMZ and RMZ of embryos following electroporation with different constructs. N, number of embryos. (D) In utero electroporation was performed at embryonic day E14.5 and analyzed 2 days later; Inhibition of Fbxo45 did not affect cell division (Ki67), apical (Sox2) or basal (Tbr2) progenitor cells, neuronal commitment (Tuj1), cell survival (cleaved Caspase-3) or the integrity of radial glia fibers (Nestin) (E) NCadHA and NC123HA were expressed at similar level by in utero electroporation. Sections of embryo brains electroporated with plasmids expressing Cherry fluorescent protein (red) and either NCad-HA (upper) or NC123-HA (lower) were visualized with anti-HA antibody (green). Scale bars: (A and B): 100 μm. (C) 50 µm for Ki67, Sox2, Tbr2, Tuj1; 100 µm for Caspase-3 and Nestin.

To test whether Fbxo45 binding to the extracellular domain of NCad may regulate neuron migration, we asked whether NC123 could overcome the migration delay caused by Rap1GAP, which inhibits NCad upregulation on the neuron surface (49). Embryos were electroporated in utero at E14.5 to express GFP, Rap1GAP and vector, NCad-HA or NC123-HA. Three days later, embryos were euthanized and the positions of the GFP-expressing neurons were visualized. As shown in Figure 7B and quantified in Figure 7C, Rap1GAP delayed neuron entry into the radial migration zone, and this delay was rescued by over-expressing NCad, as expected (P=1.3×10^−6^). However, NC123 did not rescue migration, suggesting that the NCad Fbxo45-binding site is required (P=0.41). To confirm that NC123 was expressed in this experiment, sections were stained for the HA tag on NCad-HA and NC123-HA. The wildtype and mutant proteins were expressed at similar levels (Figure 7E). These results suggest that NCad interaction with Fbxo45 or other proteins through its SPRY motifs is required to neurons to migrate into the cortical plate from the multipolar zone.

## DISCUSSION

NCad plays important roles during brain development, including stabilizing the ventricular zone, regulating neuron migration, and coordinating axonal, dendritic and synaptic differentiation (6, 7, 11-13, 22, 49). While NCad-dependent cell-cell homophilic adhesion is presumably important for many of these processes, we recently found that NCad-dependent migration into the cortical plate does not require NCad homophilic adhesion (30). Our new results suggest that NCad may need Fbxo45 to trigger migration into the cortical plate. Fbxo45 appears to be secreted by an unconventional pathway, circumventing the usual ER to Golgi route, and binds through its SPRY domain to SPRY consensus motifs in NCad EC1. These motifs are required for NCad to support neuron migration into the cortical plate but are dispensable for cell-cell adhesion. While these results do not exclude the possibility that another protein binds the same residues in NCad as Fbxo45, a parsimonious explanation is that neuron migration into the cortical plate is stimulated when Fbxo45 is secreted and binds NCad.

A previous study detected NCad binding to Fbxo45 and proposed that the interaction occurred inside the cell (34). However, the specific mechanism was not addressed. Our results suggest that Fbxo45 is actively secreted from cells by a non-classical, signal peptide-independent mechanism that is insensitive to BFA. Several other proteins undergo unconventional secretion, including cytokine interleukin-1 (50), thioredoxin (51), fibroblast growth factors FGF-1 (52) and FGF-2 (53), α-Synuclein (54, 55), Galectins (56), phosphoglucose isomerase/autocrine motility factor (57) and heat shock proteins (47). However, the mechanisms of unconventional secretion are still unclear. The lack of conserved signals, relatively small amounts of proteins secreted, and apparent variety of mechanisms have presented technical challenges (45, 47, 58). One mechanism, known as autophagy-mediated secretion, involves chaperone-mediated autophagy (CMA) to translocate proteins into lysosomes, followed by lysosome fusion with the plasma membrane (59). A loose consensus for CMA targeting contains a glutamine preceded or followed by four residues including at least one basic (K or R), one acidic (D or E), and one hydrophobic (I, L, V or F) residue. Fbxo45 contains such a sequence (residues 225-229: QIGER). However, deleting this sequence did not reproducibly inhibit Fbxo45 secretion (data not shown). It is possible that Fbxo45 secretion involves CMA and that variability in secretion may stem from variable levels of autophagy between experiments.

Mutational analysis of the Fbxo45-NCad interaction suggests that the Fbxo45 SPRY domain makes canonical interactions with three candidate SPRY motifs in NCad EC1-2 ([DE][IL]NXN (39)) that are not conserved in ECad (Figure 3 and 4). We found that swapping NCad motif 1 with the corresponding residues of ECad strongly inhibits Fbxo45 binding. This motif lies on the opposite surface from the EC1-EC1 interface in the strand-swap NCad dimer (41), so is unlikely to regulate NCad trans dimerization between cells. However, this surface has been implicated in cis interactions between adjacent cadherins on the same membrane (41).

Mutations here cause measurable changes in mechanical coupling to the actin cytoskeleton but do not affect cell-cell adhesion (60). The same region of ECad is important for a proposed conformational change involved in inside-out activation (61, 62). Mutation of a nearby residue in ECad is found in some cancers and also inhibits inside-out activation (62). Therefore, trans interactions with Fbxo45 may affect cis interactions between NCad molecules on the same cell or modulate NCad conformational activation.

Previous studies showed that interfering with NCad surface expression by inhibiting Reelin, Rap1, Rab proteins and Drebin-like, causes a delay in neuron entry into the cortical plate (22, 23, 49). We found that inhibiting Fbxo45 expression with shRNA caused a similar delay in entry into the cortical plate (Figure 7A and C). There appeared to be no significant effects on neurogenesis, cell differentiation, apoptosis or local cytoarchitecture. This suggests that Fbxo45 expression is needed at approximately the same stage as NCad. We were unable to identify mutations in Fbxo45 that inhibit secretion, so were unable to directly test whether Fbxo45 secretion is important in vivo. Instead, we performed an indirect experiment, testing whether Fbxo45-binding sites are required for over-expressed NCad to overcome the effect of Rap1GAP, which reduces NCad surface levels and inhibits migration (49). Forced expression of exogenous NCad could rescue the migration defect while the Nc123 mutant could not (Figure 7B and C). Therefore, the NCad Fbxo45-binding sites were required, consistent with binding of Fbxo45 (or another unidentified protein) to NCad before neurons enter the cortical plate.

It remains a puzzle that Fbxo45 has intracellular and extracellular functions. Inside the cell, Fbxo45 binds MycBP2 and Skp1 to stimulate ubiquitylation and turnover of Par-4, mTOR, Munc-13 and p73 (33, 63-65). This intracellular function is thought to regulate the epithelial-mesenchymal transition and synaptic function (33, 66). However, our results suggest that Fbxo45 is also secreted and acts outside the cell. Other unconventionally secreted proteins have different functions inside and outside the cell. Heat shock protein 70 (Hsp70) is an intracellular chaperone, but stress stimulates Hsp70 secretion and extracellular Hsp70 activates macrophages (47, 67). Vasohibins catalyze tubulin detyrosination but when secreted regulate angiogenesis (68, 69). As another example, phosphoglucose isomerase is a key metabolic enzyme, but when secreted it is known as autocrine motility factor and stimulates cell migration by binding to specific cell surface receptors (57). By analogy, we can hypothesize that Fbxo45 is secreted under specific biological conditions and that secreted Fbxo45 regulates NCad non-adhesive functions.

In conclusion, Fbxo45 appears to be secreted by a non-classical mechanism. Outside the cell, it binds to SPRY motifs in EC1-2 of NCad. Mutating these motifs inhibits NCad function in neuron migration without affecting cell-cell adhesion. Therefore, secreted Fbxo45, or a protein with overlapping binding requirements, likely regulates NCad during neuron migration. This suggests that Fbxo45 has different functions depending on whether it is intra- and extra-cellular. More studies are warranted to characterize the mechanism and function of the secreted form.

## MATERIALS AND METHODS

### BioID and expression plasmid cloning

To generate N5-BirA*, a PCR product of myc-BirA* (35700; Addgene) was ligated into XhoI and SpeI cleaved pCAG-mNCad-HA, which contains XhoI and SpeI sites introduced into the region between EC5 and the transmembrane domain (49). GFP-NCad contains GFP inserted at the same position. For N2-BirA* cloning, PCR products of myc-BirA* and NCad EC2 were sequentially ligated to pBluescript II KS (+) and the final insert was ligated to pCAG-NCad-HA using EcoRI and KpnI. E5-BirA* was created by ligating a PCR product of murine ECad EC1-5 to the transmembrane and intracellular regions of N5-BirA* using EcoRI and XhoI. The murine ECad template was a kind gift of Masatoshi Takeichi (70). pCAGGFP (Addgene) and pCAGCherryFP were used for co-transfection. The NC1, NC2, NC3, NC12 and NC123 mutations were created using standard site-directed mutagenesis using Kapa HiFi DNA polymerase (Kapa Biosystems) according to manufacturer’s instructions. Murine Fbxo45 (Harvard Plasmid Database) was inserted to pCAG vector possessing T7 tag using BamHI and NotI. Fbxo45 deletion mutants were generated by site-directed mutagenesis. All constructs were confirmed by sequencing. pLKO.1 Fbxo45 shRNA plasmids shFbxo45b (TRCN0000201180, target sequence GACATGGAGGATAAGACTTTA) and shFbxo45a (TRCN0000339817 target sequence TGGAATCTGGTGGACAATAAT) were obtained from Sigma. When co-transfected with Flag-Fbxo45, these shRNAs inhibited expression by 58 % and 53 % respectively.

### Cell culture and transfection

HEK293, HeLa and CHO-K1 cells were maintained in Dulbecco’s modified Eagle’s medium (DMEM) with 10 % fetal bovine serum (FBS) and penicillin/streptomycin (100 U/ml) and transfected using Lipofectamine 2000 (Thermo Fisher Scientific) according to the manufacturer’s instructions. Unless otherwise stated, transfected cells were incubated 40-48 hours before protein analysis or fixation for immunofluorescence. Cortical neurons from embryonic day 18 (E18) Sprague–Dawley rats (Charles River) or E15.5-16.5 CD-1 mice (Charles River) from were prepared as reported previously (71) and maintained in Neurobasal media with B-27 supplements (Thermo Fisher Scientific). All animal procedures were approved by the Fred Hutchinson Cancer Research Center Institutional Animal Care and Use Committee. Neurons were transfected using Lipofectamine 2000. Hippocampal neurons were prepared as reported previously (71), transfected after 3 or 5 days in vitro (DIV), and fixed 2 days later.

### HEK293-neuron trans-biotinylation (BioID)

HEK293 cells were transfected with pCAG plasmids encoding GFP (experiment 1), E5-BirA* (experiment 2), N5-BirA* or N2-BirA* (both experiments). After 24 hours, transfected cells were detached with 5 mM EDTA in PBS, centrifuged at 1,000 rpm for 5 minutes, suspended in Neurobasal media with 10 % horse serum, penicillin/streptomycin (100 U/ml) and B-27 supplements, and added to rat cortical primary cultures that had been prepared 3 days previously. About 10-18 × 10^6^ HEK293 cells were added to 100 × 10^6^ rat cortical neurons per sample. After overnight incubation, media were changed to fresh Neurobasal media containing 5 % Horse Serum, penicillin/streptomycin (100 U/ml), B-27 supplements, 50 μM biotin and 1 mM ATP. After 24 hours, mixed cells were washed with cold PBS two times and harvested in 1 % Triton X-100 in PBS buffer with 1 mM EDTA and Complete protease inhibitors (Roche). The lysates were centrifuged at 14,000 rpm for 15 minutes and the supernatant mixed with Strepavidin Sepharose High Performance (Sigma-Aldrich) and rotated for 5 hours at 4 **°**C. Streptavidin beads were washed three times with Buffer 1 (20 mM Tris-HCl, pH 7.4, 0.1 % NP-40, 1 M NaCl, 5 mM EDTA, pH 8.0) followed by Buffer 2 (2 mM Tris-HCL, pH 7.4, 0.1 % NP-40, 0.5 mM EDTA) and Buffer 3 (1 M urea, 10 mM Tris-HCl, pH 7.4, 0.1 % NP-40, 1 mM EDTA), 2 times each. A final wash was performed with 50 mM Tris-HCl, pH 7.4 to remove urea and detergents. The beads were equilibrated with 50 mM ammonium bicarbonate, pH 8.0 and digested with trypsin overnight at 37 **°**C with gentle agitation. Additional trypsin was added for a further 2 hours at 37 **°**C. Digestion was stopped by adding formic acid to 1 % final concentration. Beads were centrifuged at 4,000 rpm for 2 minutes and the supernatant was transferred to a new tube. Beads were rinsed twice more with water and all supernatants were combined and lyophilized. The peptides were cleaned using ZipTip micro-C18 (Millipore Corporation) and analyzed by LC-MS/MS.

### LC-MS/MS analysis for BioID

LC-MS/MS analysis was performed with an Easy-nLC 1000 (Thermo Scientific) coupled to an Orbitrap Elite mass spectrometer (Thermo Scientific). The LC system, configured in a vented format, consisted of a fused-silica nanospray needle (PicoTip™ emitter, 75 µm ID, New Objective) packed in-house with Magic C18 AQ 100Å reverse-phase media (Michrom Bioresources Inc.) (25 cm), and a trap (IntegraFrit Capillary, 100 µm ID, New Objective) containing Magic C18 AQ 200Å (2 cm). The peptide sample was diluted in 10 µL of 2% acetonitrile and 0.1 % formic acid in water and 8 µL was loaded onto the column and separated using a two-mobile-phase system consisting of 0.1 % formic acid in water (A) and 0.1 % acetic acid in acetonitrile (B). A 90 minute gradient from 7 % to 35 % acetonitrile in 0.1 % formic acid at a flow rate of 400 nL/min was used for chromatographic separations. The mass spectrometer was operated in a data-dependent MS/MS mode over the m/z range of 400-1800. The mass resolution was set at 120,000. For each cycle, the 15 most abundant ions from the scan were selected for MS/MS analysis using 35 % normalized collision energy. Selected ions were dynamically excluded for 30 seconds.

### Cadherin-Fc fusion proteins

DNA sequences encoding the ectodomains of ECad and NCad^W161A^ were PCR amplified and cut with MfeI and AgeI (ECad) or EcoRI and AgeI (NCad) and cloned into a plasmid (pcDNA3.1-ApoER2-Fc (15)) that had been cut EcoRI and AgeI. The murine ECad template was a kind gift of Masatoshi Takeichi (70) and murine NCad^W161A^ template was from (30). The plasmids were confirmed by sequencing. Fusion proteins were produced in HEK293T cells. For each construct, five 6-cm plates were transfected with a total of 120 µg of DNA using the calcium phosphate method. Media were removed 6 hours later and gently replaced with serum-free DMEM. Two days later, media were harvested and concentrated using Amicon Centricon YM-100 centrifugal filters. Protein concentration and purity were estimated by SDS PAGE with a BSA standard. Approximately 250 µg of each fusion protein was mixed with 250 µL packed volume Protein A/G PLUS-Agarose (Santa Cruz Biotechnology) and unbound proteins washed away using PBS.

### Purification of proteins interacting with NCadW161A and not with ECad

Brains from 21 mouse embryos (E16.5) were homogenized with a Dounce homogenizer in a total volume of 8 mL of Buffer A (100 mM NaCl, 20 mM Hepes pH 7.5, 2 mM CaCl_2_, 0.2 mM EDTA, Complete protease inhibitors) containing 1 % Triton X-100. The homogenate was centrifuged at 10,000 x g for 30 minutes at 4 **°**C. The supernatant was adjusted to contain 0.3 µg/ml DNaseI and mixed with Protein A/G PLUS-Agarose (120 µL packed volume) in a BioRad “Biospin” mini column for 30 minutes at 4 **°**C. Unbound material was mixed with ECad-Fc bound to Protein A/G PLUS-Agarose (440 µL packed volume, estimated 220 µg ECad-Fc) for 90 minutes at 4 **°**C. Unbound material was mixed with NCad^W161A^-Fc bound to Protein A/G PLUS-Agarose (440 µL packed volume, estimated 220 µg ECad-Fc) for 2 hours at 4 **°**C. Finally, all three columns were washed eight times with 1 mL Buffer A containing 0.1 % Triton X-100, once with 1 mL of 20 mM Hepes pH 7.5, 2 mM CaCl_2_ and once with 1 mL of H_2_O before eluting twice with 1 mL of 0.1 % formic acid in H_2_O. Eluates were neutralized with 100 µL 1 M ammonium bicarbonate, frozen, and lyophilized. Lyophilized proteins were dissolved in SDS-PAGE sample buffer and resolved by SDS-PAGE (BioRad precast 4-15 % gradient gel). The gel was stained with Simple Blue Safe Stain (Invitrogen). Sections of the gel above and below the Cadherin-Fc bands were excised and analyzed by trypsin digestion and mass spectrometry. Briefly, **t**he gel pieces were destained with 25 mM ammonium bicarbonate in 50 % acetonitrile, and subsequently dehydrated using acetonitrile. The protein was digested for overnight with 5 ng/µL trypsin (Promega Corporation) in 50 mM ammonium bicarbonate at 37 °C. The peptides were extracted using 5 % v/v formic acid in water, then with acetonitrile. The pooled extracts were dried under vacuum and cleaned using ZipTip™ C18 (Millipore Corporation) before the subsequent MS analysis.

### LC-MS/MS analysis for Affinity Purification

LC-MS/MS analysis was performed with an Eksigent NanoLC-2D system (Eksigent/AB Sciex) coupled to an LTQ Orbitrap mass spectrometer (Thermo Scientific). The LC system configured in a vented format (72) consisted of a fused-silica nanospray needle (PicoTip™ emitter, 75 µm ID, New Objective) packed in-house with Magic C18 AQ 100 Å reverse-phase media (Michrom Bioresources Inc.) (25 cm), and a trap (IntegraFrit™ Capillary, 100 µm ID, New Objective) containing Magic C18 AQ 200Å (2 cm). The peptide sample was diluted in 10 µL of 2 % acetonitrile and 0.1 % formic acid in water and 8 µL was loaded onto the column and separated using a two-mobile-phase system consisting of 0.1 % formic acid in water (A) and 0.1 % acetic acid in acetonitrile (B). A 60 min gradient from 7 % to 35 % acetonitrile in 0.1 % formic acid at a flow rate of 400 nL/min was used for chromatographic separations. The mass spectrometer was operated in a data-dependent MS/MS mode over the m/z range of 400-1800. The mass resolution was set at 60,000. For each cycle, the 5 most abundant ions from the scan were selected for MS/MS analysis using 35 % normalized collision energy. Selected ions were dynamically excluded for 45 s.

### MS data analysis

Data analysis was performed using Proteome Discoverer 1.4 (Thermo Scientific). The data were searched against Uniprot human (March 19, 2014), rat (January 9, 2015), and mouse (March 19, 2014) protein databases, and against common contaminants (http://www.thegpm.org/crap/). Trypsin was set as the enzyme with maximum missed cleavages set to 2. The precursor ion tolerance was set to 10 ppm, and the fragment ion tolerance was set to 0.6 Da. SEQUEST (73) was used for search, and search results were run through Percolator (74) for scoring.

### Immunoprecipitation and pull down

Cells were washed with cold PBS two times and harvested in 1% Triton X-100 in PBS with protease/phosphatase inhibitors. The lysates were centrifuged at 14,000 rpm for 15 minutes and supernatant mixed with T7 Tag Antibody Agarose (EMD Millipore) for immunoprecipitation or NeutrAvidin (Thermo Fisher Scientific) for the streptavidin pull down for 2 hours. Agarose beads were washed three times with lysis buffer, eluded with SDS sampling buffer, and analyzed by SDS-PAGE and western blotting. The following antibodies were used for immunoprecipitation and Western blotting: mouse N-cadherin (Thermo Fisher Scientific), HRP conjugated Strepavidin (Jackson ImmunoResearch Laboratories, Inc.), mouse anti-tubulin (Santa Cruz Biotechnology, Inc.), mouse anti-T7 (EMD Millipore), mouse anti-HA (HA.11) (BioLegend), rabbit anti-HA (Bethyl, Montgomery) and mouse Flag M2 (Sigma-Aldrich). Rabbit Fbxo45 antibody was kindly provided by Kojo S. J. Elenitoba-Johnson (University of Pennsylvania).

### Immunofluorescence

HeLa cells were seeded onto collagen IV-coated 12-mm coverslips at 10^5^ cells/well of a 24-well plate and transfected immediately with 400 ng of pCAG-NCad-HA or pCAG-NC123-HA using Lipofectamine 2000. The media were changed 6 hr later and 2 days later the cells were fixed with 4 % paraformaldehyde for 15 min at room temperature. Coverslips were washed in PBS and blocked for 1 hour in 1 % BSA, 5 % normal goat serum in PBS, incubated with primary antibody for 1 hour, washed, incubated with secondary antibodies for 1 hour, washed, and mounted in Prolong Glass (Thermo Fisher Scientific). All procedures were performed at room temperature unless otherwise specified. Primary antibodies were: rabbit anti-NCad extracellular H-63 (Santa Cruz Biotechnology), mouse anti-EGFR extracellular (528) (Calbiochem), rat anti-HA 3F10 (Sigma). Secondary antibodies were: Alexa 568 goat anti-rabbit IgG (H+L), Alexa 488 goat anti-mouse IgG (H+L), Alexa 647 goat anti-rat IgG (H+L), all from Jackson Immunoresearch. No staining with HA antibody was detected, showing that the cells were not permeabilized. NCad, EGFR and HA antibodies all stained cytoplasmic vesicles if the cells were permeabilized with 0.1 % Triton X100 before blocking and antibody addition (data not shown).

### Preparation of secreted proteins and total cell lysates

To collect secreted proteins, cells were incubated in serum-free DMEM (for HeLa) or Neurobasal Media (for primary cortical neurons) for 20-24 hours at 37 **°**C with 5 % CO_2_. The conditioned medium was centrifuged at 3,000 rpm for 5 minutes to remove cell debris. Supernatants were collected and concentrated using Amicon Ultra, 10 kDa NMWL (EMD Millipore). After removing the conditioned medium, cells were washed with cold PBS and harvested in 1% Triton X-100 in PBS buffer with protease/phosphatase inhibitors. Triton-insoluble material was removed at 14,000 rpm for 15 minutes. Samples of the conditioned medium and cell lysate were analyzed by SDS polyacrylamide gel electrophoresis and Western blotting.

### Short-term aggregation assay

CHO-K1 cells were co-transfected with GFP or mCherry and indicated NCad constructs. After 24 hours, transfected cells were washed twice with pre-warmed HCMF (137 mM NaCl, 5.4 mM KCl, 0.63 mM Na_2_HPO_4_, 5.5 mM Glucose, 10 mM HEPES, pH 7.4) and placed in suspension using 0.01 % trypsin, 1 mM CaCl_2_ in HCMF. Suspended cells were centrifuged at 1,000 rpm for 5 minutes, resuspended in 0.5 % soybean trypsin inhibitor in HCMF, then washed three times with cold HCMF and counted. 200,000 cells in HCMF were added to 1 % BSA-coated 24-well plates with and without 2 mM CaCl_2_. Cells were shaken at 80 rpm for 20 minutes at 37 **°**C and images were collected using 2X or 4X objectives. Cell aggregation assays were performed three biological replicates. Data were quantified using Analyze Particles in Image J (75).

### In utero electroporation

In utero microinjection and electroporation was performed at E14.5 essentially as described (76) using timed pregnant CD-1 mice (Charles River Laboratories). In brief, mice were anesthetized and the midline incision the uterine horns were exposed. Plasmid solution was injected into the lateral ventricle using needles for injection that were pulled from Wiretrol II glass capillaries (Drummond Scientific) and calibrated for 1-μl injections. DNA solutions were mixed in 10 mM Tris, pH 8.0 with 0.01 % Fast Green. The embryo in the uterus was placed between the forceps-type electrodes (Nepagene) with 5-mm pads and electroporated with five 50-ms pulses of 45 V using the ECM830 electroporation system (Harvard Apparatus). The uterine horns were then placed back into the abdominal cavity to continue normal brain development. Three days later, mice were euthanized and embryo brains were sectioned and GFP visualized.

Immunohistochemistry to detect HA-tagged NCad was performed essentially as described (49). The following additional antibodies were used: Ki67 (BD Biosciences 556003), Sox2 (Cell Signaling Technology 4900), Tbr2 (Abcam ab23345), Tuj1 (Covance MMS-435P), Cleaved Caspase3 (Cell Signaling Technology 9661), Nestin (Millipore MAB353).

### Statistical analysis

Statistical analysis was performed with GraphPad Prism 7.0 (La Jolla, CA). Statistical significance was determined by two-tailed unpaired t-test. Data are reported as mean±SEM and the statistical significance was set as p<0.05.

## AUTHOR CONTRIBUTIONS

Y.N., E.C.J., E.K, H.C. and Y.J. performed experiments with technical assistance from J.A.C. Y.N. and J.A.C. wrote and revised the manuscript with input from Y.J.

## ACKNOWLEDGEMENTS

We are very grateful to Barry Gumbiner and Kojo S. J. Elenitoba-Johnson for helpful discussions and sharing reagents. We thank staff of the Fred Hutch shared resources, especially Phil Gafken, Yuko Ogata, David McDonald, Lena Schroeder and Julio Vazquez, for proteomics and imaging; Alexander (Sasha) Straight for technical assistance, and members of the Cooper laboratory for encouragement. This work was supported by grants R01 NS080194 and GM109463 from the U.S. Public Health Service and grants J.0129.15, J.0179.16 and T.0243.18 from the FNRS. FHCRC imaging and proteomics laboratories are supported by P01 CA015704.

