## Supplemental tables for "Fbxo45 binds SPRY motifs in the extracellular domain of N-cadherin and regulates neuron migration during brain development" (from another version of this preprint: version 2, 2020-03-31)

**Supplementary Table 1. Rat proteins detected by BioID with NCad-BirA*.**

Rat proteins detected by LC-MS/MS from two independent experiments. The numbers represent the score and the data was sorted by the score of N2-BirA* in experiment 2 as a descending manner. The score is the sum of all peptide Xcorr values above the specified score threshold. The score threshold is calculated as follows: 0.8 + peptide charge X peptide relevance factor. 0.4 was used for the peptide relevance factor. For each spectrum, only the highest-scoring match is used. Blank cells indicate that no peptide was detected in the sample.

**Supplementary Table 2. Human proteins detected by BioID with NCad-BirA*.**

Human proteins detected by mass spectrometry from the same two experiments as Table 1.

**Supplementary Table 3. Mouse proteins detected by affinity purification as a binding candidate of NCad EC W161A.**

Proteins from gel regions above and below of the cadherin-Fc bands in Figure 1F were digested with trypsin and analyzed by LC-MS/MS. The numbers represent the score calculated as described in Table 1. For each spectrum, only the highest-scoring match is used. Blank cells indicate that no peptide was detected in the sample.

### Supplementary Table 1

| Accession | Rat Gene | Protein description | Experiment 1 | | | Experiment 2 | | |
|---|---|---|---|---|---|---|---|---|
| | | | GFP | N5-BirA* | N2-BirA* | E5-BirA* | N5-BirA* | N2-BirA* |
| D4A2D3 | Mycbp2 | Protein Mycbp2 (Fragment) | - | 26.15 | 49.91 | - | 26.88 | 46.08 |
| F1M8Q1 | Fbxo45 | F-box/SPRY domain-containing protein 1 (Fragment) | - | 2.99 | 8.48 | - | 2.99 | 8.65 |
| F1LYJ8 | Gramd4 | Protein Gramd4 (Fragment) | - | 2.93 | 3.10 | - | - | 6.01 |
| D3ZQN7 | Lamb1 | Protein Lamb1 | - | - | 8.00 | - | - | 5.51 |
| F1LRM0 | Scaper | Protein Scaper | - | - | 2.13 | - | 2.51 | 5.50 |
| P54777 | Pex6 | Peroxisome assembly factor 2 | - | - | 8.77 | - | - | 5.19 |
| Q63362 | Ndufa5 | NADH dehydrogenase [ubiquinone] 1 alpha subcomplex subunit 5 | - | 2.67 | - | - | - | 3.38 |
| O35796 | C1qbp | Complement component 1 Q subcomponent-binding protein, mitochondrial | - | 3.58 | - | - | 3.13 | 3.32 |
| M0RB15 | Rpp14 | Uncharacterized protein | - | - | 2.94 | - | 4.36 | 3.16 |
| D3ZPE6 | Mrpl51 | Mitochondrial ribosomal protein L51 (Predicted) | - | - | 3.47 | - | 3.34 | 2.94 |
| B2GV24 | Ufl1 | E3 UFM1-protein ligase 1 | - | 2.48 | - | - | 2.19 | 2.93 |
| F1LZW2 | Dgkk | Diacylglycerol kinase (Fragment) | - | 2.72 | - | - | 3.15 | 2.83 |
| P36972 | Aprt | Adenine phosphoribosyltransferase | - | 3.38 | - | - | - | 2.69 |
| P11951 | Cox6c2 | Cytochrome c oxidase subunit 6C-2 | - | 3.15 | 3.44 | - | - | 2.66 |
| Q6P684 | Smn1 | Survival motor neuron 1 | - | 2.13 | - | - | - | 2.50 |
| D3ZFF4 | Lclat1 | Protein Lclat1 | - | - | 6.21 | - | 1.82 | 2.46 |
| F1LQE9 | Ppp1r10 | Uncharacterized protein | - | - | 3.88 | - | - | 2.43 |
| B2RZD2 | Tmem126b | Complex I assembly factor TMEM126B, mitochondrial | - | 2.32 | - | - | 3.18 | 2.27 |
| Q3T1K9 | Nicn1 | Nicolin 1 | - | - | 3.18 | - | 0.00 | 2.23 |
| Q6AYA1 | Gar1 | H/ACA ribonucleoprotein complex subunit 1 | - | 2.20 | 2.06 | - | 2.46 | 2.22 |
| Q5BJS0 | Dhx30 | Putative ATP-dependent RNA helicase DHX30 | - | - | 2.41 | - | 2.04 | 2.11 |
| D3ZI07 | Kif3b | Kinesin-like protein | - | 2.69 | 2.97 | - | 3.19 | 2.00 |
| P50137 | Tkt | Transketolase | - | 10.03 | 7.29 | - | 2.59 | - |
| D4ABT8 | Hnrnpul2 | Protein Hnrnpul2 | - | 7.52 | 7.17 | - | 2.01 | - |
| M0R7U6 | Sowahc | Protein Sowahc | - | 3.16 | 3.90 | - | 2.72 | - |
| F1MAF2 | Akap17a | Protein Akap17a | - | 3.17 | 3.29 | - | 3.59 | - |
| D3ZHV9 | Misp | Hypothetical LOC500797, isoform CRA_a | - | 2.14 | 2.03 | - | 2.94 | - |
| Q07009 | Capn2 | Calpain-2 catalytic subunit | - | 3.10 | - | - | 2.65 | - |
| D3ZZQ6 | Ints4 | Protein Ints4 | - | 2.18 | - | - | 2.53 | - |

### Supplementary Table 2

| Accession | Human Gene | Description | Experiment 1 | | | Experiment 2 | | |
|---|---|---|---|---|---|---|---|---|
| | | | GFP | N5-BirA* | N2-BirA* | E5-BirA* | N5-BirA* | N2-BirA* |
| O75592-2 | MYCBP2 | Isoform 2 of Probable E3 ubiquitin-protein ligase MYCBP2 | - | 47.55 | 127.74 | - | 70.14 | 112.99 |
| Q13751 | LAMB3 | Laminin subunit beta-3 | - | - | 2.71 | - | - | 8.56 |
| Q9BR76 | CORO1B | Coronin-1B | - | 3.81 | 8.88 | - | 4.30 | 7.33 |
| P46108-2 | CRK | Isoform Crk-I of Adapter molecule crk | - | - | 3.31 | - | 2.33 | 7.14 |
| A6NJ11 | UFD1L | Ubiquitin fusion degradation protein 1 homolog | - | 3.12 | 3.33 | - | 8.55 | 5.86 |
| Q9NT62-2 | ATG3 | Isoform 2 of Ubiquitin-like-conjugating enzyme ATG3 | - | - | 2.13 | - | - | 5.22 |
| Q9Y3A6 | TMED5 | Transmembrane emp24 domain-containing protein 5 | - | 3.37 | - | - | 3.60 | 5.19 |
| Q5SSJ5-3 | HP1BP3 | Isoform 3 of Heterochromatin protein 1-binding protein 3 | - | 2.96 | 3.59 | - | 0.00 | 5.16 |
| Q15750-2 | TAB1 | Isoform 2 of TGF-beta-activated kinase 1 and MAP3K7-binding protein 1 | - | - | 3.50 | - | 2.15 | 4.38 |
| B4DEK4 | SNX2 | Sorting nexin-2 | - | 4.88 | 2.84 | - | 5.53 | 4.25 |
| Q8TCG1-2 | KIAA1524 | Isoform 2 of Protein CIP2A | - | 3.70 | 4.30 | - | - | 3.65 |
| Q09472 | EP300 | Histone acetyltransferase p300 | - | 3.18 | 2.44 | - | - | 3.34 |
| Q9UHI6 | DDX20 | Probable ATP-dependent RNA helicase DDX20 | - | - | 2.52 | - | 3.70 | 3.18 |
| O95400 | CD2BP2 | CD2 antigen cytoplasmic tail-binding protein 2 | - | 3.61 | 6.02 | - | 3.20 | 3.07 |
| Q9UHG3 | PCYOX1 | Prenylcysteine oxidase 1 | - | - | 5.62 | - | - | 2.80 |
| P11047 | LAMC1 | Laminin subunit gamma-1 | - | - | 22.15 | - | - | 2.80 |
| J3KRS1 | DHRS7B | Dehydrogenase/reductase SDR family member 7B (Fragment) | - | 2.78 | 3.79 | - | - | 2.78 |
| O15063 | KIAA0355 | Uncharacterized protein KIAA0355 | - | 2.90 | 2.63 | - | 5.67 | 2.77 |
| P67775-2 | PPP2CA | Isoform 2 of Serine/threonine-protein phosphatase 2A catalytic subunit alpha isoform | - | 1.66 | 1.90 | - | - | 2.70 |
| B4DT16 | BCL11A | B-cell lymphoma/leukemia 11A | - | 2.37 | - | - | - | 2.69 |
| E7ERI3 | TARS | Threonine--tRNA ligase, cytoplasmic | - | 1.92 | - | - | 2.73 | 2.63 |
| Q70E73-5 | RAPH1 | Isoform RMO1c of Ras-associated and pleckstrin homology domains-containing protein 1 | - | - | 4.16 | - | - | 2.60 |
| O76024 | WFS1 | Wolframin | - | - | 2.60 | - | 2.29 | 2.51 |
| D6REK3 | CWC27 | Peptidyl-prolyl cis-trans isomerase CWC27 homolog | - | - | 2.51 | - | - | 2.50 |
| O14617-4 | AP3D1 | Isoform 4 of AP-3 complex subunit delta-1 | - | - | 0.00 | - | 2.61 | 2.46 |
| Q99848 | EBNA1BP2 | Probable rRNA-processing protein EBP2 | - | - | 1.94 | - | 1.83 | 2.44 |
| H0Y380 | PTPRF | Receptor-type tyrosine-protein phosphatase F (Fragment) | - | - | 2.38 | - | 2.07 | 2.16 |
| Q9UL03-3 | INTS6 | Isoform 3 of Integrator complex subunit 6 | - | 2.47 | 3.12 | - | - | 2.15 |
| O75382-4 | TRIM3 | Isoform 4 of Tripartite motif-containing protein 3 | - | - | 3.98 | - | - | 2.12 |
| M0QYC5 | SIN3B | Paired amphipathic helix protein Sin3b | - | - | 3.68 | - | 3.63 | 2.08 |
| H3BPL5 | CCDC101 | SAGA-associated factor 29 homolog | - | - | 2.50 | - | 2.06 | 2.06 |
| Q9H944 | MED20 | Mediator of RNA polymerase II transcription subunit 20 | - | - | 3.32 | - | - | 2.03 |
| Q8ND04 | SMG8 | Protein SMG8 | - | 4.56 | - | - | - | 2.01 |
| Q9H061 | TMEM126A | Transmembrane protein 126A | - | 2.76 | - | - | 3.52 | - |
| P08621-2 | SNRNP70 | Isoform 2 of U1 small nuclear ribonucleoprotein 70 kDa | - | 5.10 | 3.18 | - | 1.79 | - |
| J3KQN6 | PPP2R5E | Serine/threonine-protein phosphatase 2A 56 kDa regulatory subunit epsilon isoform | - | 4.01 | 4.50 | - | 2.38 | - |
| E9PGB5 | SFXN4 | Sideroflexin-4 | - | 2.02 | 1.64 | - | 1.72 | - |
| Q0PRL4 | FOXP2 | Forkhead box P2 variant 3 | - | 3.39 | - | - | 3.02 | - |
| H7C0X5 | CPVL | Probable serine carboxypeptidase CPVL (Fragment) | - | 2.39 | 2.23 | - | 2.07 | - |
| E9PM04 | DCUN1D5 | DCN1-like protein | - | 3.43 | - | - | 2.87 | - |
| F5GXS7 | METAP1 | Methionine aminopeptidase | - | 2.80 | 4.69 | - | 2.07 | - |
| F5H6N1 | MAGOHB | Protein mago nashi homolog 2 | - | 3.09 | 3.03 | - | 2.66 | - |
| K7EKI4 | MRPL4 | 39S ribosomal protein L4, mitochondrial | - | 2.48 | - | - | 2.09 | - |
| S4R432 | CBWD3 | COBW domain-containing protein 3 (Fragment) | - | 7.27 | - | - | 1.99 | - |
| Q5H937 | PKIG | cAMP-dependent protein kinase inhibitor gamma | - | - | 3.10 | - | 1.89 | - |
| E7EN38 | RAD50 | DNA repair protein RAD50 (Fragment) | - | - | 3.47 | - | 3.78 | - |
| Q7Z2Z2-2 | EFTUD1 | Isoform 2 of Elongation factor Tu GTP-binding domain-containing protein 1 | - | - | 2.81 | - | 1.85 | - |
| B3KT28 | FAF1 | FAS-associated factor 1 | - | - | 2.27 | - | 2.29 | - |
| B4DVY7 | FLOT1 | Flotillin-1 | - | - | 4.30 | - | 2.58 | - |
| D6RBM3 | CASP6 | Caspase-6 subunit p18 | - | - | 3.13 | - | 2.75 | - |
| P09496-5 | CLTA | Isoform 5 of Clathrin light chain A | - | - | 7.16 | - | 4.60 | - |
| F5GX19 | LAMTOR1 | Ragulator complex protein LAMTOR1 | - | - | 2.70 | - | 2.55 | - |

### Supplementary Table 3

| Accession | Gene | Description | Ecad-Fc | Ncad$^{W161A}$-Fc |
|---|---|---|---|---|
| Q7TPH6-2 | Mycbp2 | Isoform 2 of Probable E3 ubiquitin-protein ligase MYCBP2 | - | 205.37 |
| Q8K3B1 | Fbxo45 | F-box/SPRY domain-containing protein 1 | - | 27.50 |
| Q8R1J9 | Tor2a | Torsin-2A | - | 21.99 |
| Q8C570 | Rae1 | mRNA export factor | - | 13.51 |
| O88838 | Spsb2 | SPRY domain-containing SOCS box protein 2 | - | 12.70 |
| P83940 | Tceb1 | Transcription elongation factor B polypeptide 1 | - | 10.66 |
| Q9D051 | Pdhb | Pyruvate dehydrogenase E1 component subunit beta, mitochondrial | - | 9.71 |
| Q9WTX5 | Skp1 | S-phase kinase-associated protein 1 | - | 7.54 |
| Q9DCL9 | Paics | Multifunctional protein ADE2 | - | 7.10 |
| A2A5N1 | Ywhab | 14-3-3 protein beta/alpha (Fragment) | - | 6.49 |
| Q9CZU6 | Cs | Citrate synthase, mitochondrial | - | 6.22 |
| O08573-2 | Lgals9 | Isoform Short of Galectin-9 | - | 5.36 |
| E9Q9B3 | Spryd3 | Protein Spryd3 | - | 5.31 |
| Q61830 | Mrc1 | Macrophage mannose receptor 1 | - | 5.23 |
| Q61753 | Phgdh | D-3-phosphoglycerate dehydrogenase | - | 4.98 |
| P11983-2 | Tcp1 | Isoform 2 of T-complex protein 1 subunit alpha | - | 4.90 |
| Q8BM88 | Ctso | Cathepsin O | - | 4.48 |
| Q6P5E4 | Uggt1 | UDP-glucose:glycoprotein glucosyltransferase 1 | - | 3.69 |
| F8WJ41 | Rps15a | 40S ribosomal protein S15a (Fragment) | - | 3.56 |
| H3BJB6 | Cct8 | T-complex protein 1 subunit theta (Fragment) | - | 2.99 |
| P59708 | Sf3b14 | Pre-mRNA branch site protein p14 | - | 2.97 |
| Q9JJV2-3 | Pfn2 | Isoform 3 of Profilin-2 | - | 2.71 |
| G3UXF3 | Cct4 | T-complex protein 1 subunit delta | - | 2.70 |
| P56379 | Mp68 | 6.8 kDa mitochondrial proteolipid | - | 2.56 |
| G3UZJ2 | Map2 | Microtubule-associated protein (Fragment) | - | 2.53 |
| H7BX22 | Ranbp1 | Ran-specific GTPase-activating protein | - | 2.52 |
| P99027 | Rplp2 | 60S acidic ribosomal protein P2 | - | 2.51 |
| D3YVF4 | Rps14 | 40S ribosomal protein S14 (Fragment) | - | 2.43 |
| Q99J74 | Tusc3 | Tumor suppressor candidate 3 | - | 2.25 |
| P55284 | Cdh5 | Cadherin-5 | - | 2.15 |
| E0CZA1 | Cct5 | T-complex protein 1 subunit epsilon (Fragment) | - | 2.14 |
| E9Q0W8 | Snrpe | Small nuclear ribonucleoprotein E | - | 2.09 |
| Q9CZM3 | Tmem33 | Transmembrane protein 33 | - | 2.07 |
| Q9Z0F5 | Ch25h | Cholesterol 25-hydroxylase | - | 2.06 |
| Q60692 | Psmb6 | Proteasome subunit beta type-6 | - | 2.01 |
| P00493 | Hprt1 | Hypoxanthine-guanine phosphoribosyltransferase | - | 2.00 |
| P62077 | Timm8b | Mitochondrial import inner membrane translocase subunit Tim8 B | - | 1.99 |
| Q07646-2 | Mest | Isoform 2 of Mesoderm-specific transcript protein | - | 1.96 |
| Q3UEG7 | Itih2 | Inter-alpha-trypsin inhibitor heavy chain H2 (Fragment) | - | 1.84 |
| Q9DCC4 | Pycrl | Pyrroline-5-carboxylate reductase 3 | - | 1.80 |
| E9Q3T0 | Gm10073 | Uncharacterized protein | - | 1.78 |
| P01029 | C4b | Complement C4-B | - | 1.77 |
| Q6P5D8 | Smchd1 | Structural maintenance of chromosomes flexible hinge domain-containing protein 1 | - | 1.73 |
| A2A7B3 | Spsb1 | SPRY domain-containing SOCS box protein 1 (Fragment) | - | 1.73 |
| Q91V01 | Lpcat3 | Lysophospholipid acyltransferase 5 | - | 1.70 |
| Q6P069-2 | Sri | Isoform 2 of Sorcin | - | 1.69 |
| Q3V1Z5 | Rps4l | Protein Rps4l | - | 1.60 |
